# DeepResearch^Eco^: A Recursive Agentic Workflow for Complex Scientific Question Answering in Ecology

**DOI:** 10.1101/2025.07.14.664755

**Authors:** Jennifer D’Souza, Endres Keno Sander, Andrei Aioanei

**Author notes:** Corresponding author. (J. D’Souza). All authors contributed equally. (E. K. Sander); (A. Aioanei). https://sites.google.com/view/jen-web/home (J. D’Souza).

## Abstract

We introduce **DeepResearch**^Eco^, a novel agentic LLM-based system for automated scientific synthesis that supports recursive, depth- and breadth-controlled exploration of original research questions—enhancing search diversity and nuance in the retrieval of relevant scientific literature. Unlike conventional retrieval-augmented generation pipelines, DeepResearch enables user-controllable synthesis with transparent reasoning and parameter-driven configurability, facilitating high-throughput integration of domain-specific evidence while maintaining analytical rigor. Applied to 49 ecological research questions, DeepResearch achieves up to a 21-fold increase in source integration and a 14.9-fold rise in sources integrated per 1,000 words. High-parameter settings yield expert-level analytical depth and contextual diversity.

Source code available at: https://github.com/sciknoworg/deep-research.

## 1. Introduction

Science requires extreme attention to detail, and large language models (LLMs) can overlook or misuse details when faced with challenging reasoning problems [1, 2]. Ensuring factual accuracy in LLM-generated content has thus become a key challenge. The current paradigm for eliciting factually grounded responses from LLMs is to use retrieval-augmented generation (RAG) [3, 4], which supplements the model’s knowledge with relevant documents from external sources. By leveraging retrieval, such agentic pipelines can explore scientific literature at a much higher throughput than human scientists—enabling comprehensive surveys that were previously impractical. However, scaling up literature exploration in this manner raises new questions about how to balance **breadth** (covering many sources) versus **depth** (deeply analyzing the evidence from each source) to produce high-quality scientific syntheses.

In this work, we introduce **DeepResearch**^Eco^, an agentic LLM-based system for complex scientific question answering and literature synthesis, which in this work is tested against research questions in the ecological sciences. DeepResearch employs a recursive retrieval and generation loop guided by explicit, user-controllable depth and breadth parameters. This design enables iterative broad exploration of the topic followed by targeted deep dives, effectively marrying a wide-ranging literature survey with in-depth analysis. Unlike prior feed-forward pipelines, our approach surfaces intermediate reasoning steps (e.g., search subqueries and extracted “learnings”) and uses them to refine subsequent queries, yielding a transparent and traceable knowledge workflow. We integrate two variants of LLM reasoning models within this framework to assess the robustness and generality of the generated research reports across different model capabilities. The result is a flexible methodology that can be tuned to either quickly scan numerous publications or rigorously drill down into specific evidence, all within an automated agentic workflow.

Specifically, we explore the following empirically driven research questions in this work: ***RQ1:*** *How similar are the reports generated by DeepResearch across different depth and breadth settings, using two variants of reasoning models, when evaluated with ROUGE (word-based) and embedding-based semantic similarity metrics?* ***RQ2:*** *How do depth and breadth parameters in automated research systems affect the quality and diversity of synthesized scientific knowledge in ecology?* ***RQ3:*** *Can high-parameter configurations in LLM-based systems achieve domain-specific synthesis capabilities that match or exceed expert-level integration, especially in ecological research contexts?* To answer these questions, we conduct extensive experiments using DeepResearch on ecological science problems, analyzing both quantitative metrics and qualitative aspects of the generated outputs. In summary, our contributions are threefold: (1) we present a novel recursive, breadth-vs-depth controllable LLM workflow for automated scientific literature review; (2) we provide an in-depth evaluation of how exploration depth and breadth impact the quality and diversity of knowledge synthesis (finding, for example, that a high-depth configuration can automatically integrate information from 111 sources—nearly 6× more than a shallow setting—and increase coverage of key concepts by 25%); and (3) we demonstrate that carefully configured, high-parameter runs can approach expert-level integration in ecology, achieving an order-of-magnitude higher information density in outputs without loss of rigor or specificity. All code and data are released under an open-source MIT license^1^ to facilitate reproducibility and future research.

## 2. Related Work

Recent developments in LLMs have led to the emergence of agentic workflows for scientific question answering and synthesis. This section reviews key systems that shape the landscape of LLM-based scientific discovery, covering agentic pipelines, human-aligned synthesis frameworks, and multi-agent reasoning systems.

### Scientific Search and Synthesis

A growing line of work focuses on using LLMs to automate scientific search and synthesis, combining document retrieval with intelligent summarization and reasoning capabilities.

PaperQA [5] introduced a modular agentic pipeline for LLM-assisted scientific question answering. It begins by retrieving relevant papers through Google Scholar using keyword and year-range queries, then constructs an embedding-based chunk database. For each user question, relevant text chunks are retrieved using maximal marginal relevance. These chunks are then summarized or marked as irrelevant, helping mitigate semantic noise and parsing errors. Finally, an LLM generates a response, using its own knowledge and optionally the summarized content. PaperQA2 [1] builds on this model by introducing a full-fledged multi-agent framework. Retrieval and generation are separated into distinct agents: a paper search agent reformulates the user query, fetches PDFs, and converts them to text; a citation traversal agent expands the corpus through citation networks; a gather-evidence agent retrieves and summarizes text chunks via dense retrieval, reranking, and contextual summarization; and a generation agent synthesizes answers from the top-ranked evidence. PaperQA2 also powers use cases beyond question answering, including Wikipedia-style summarization (WikiCrow) and contradiction detection (ContraCrow), the latter benchmarking whether scientific claims contradict prior literature.

ORKG Ask [6] offers a complementary approach rooted in scholarly infrastructure. It combines semantic search over a 70+ million article index (from CORE [7]) with knowledge extraction via LLMs. The search interface returns top-ranked articles using vector similarity (via Nomic embeddings), and the LLM generates a synthesis of the top 5 results. This is augmented by LLMs4Synthesis [8], a framework that structures synthesis tasks into paper-wise, methodological, and thematic categories. Syntheses are generated using structured prompts and evaluated using GPT-4 as an LLM-as-a-judge. This highlights an important emerging research direction: leveraging LLM-as-a-judge [9] as a scalable and effective approach for evaluating scientific tasks. RLAIF (reinforcement learning with AI feedback) is further applied to optimize open-source models (e.g., Mistral-7B) for factuality and clarity.

### Iterative, Structured, and Human-Aligned Research Workflows

Beyond retrieval and synthe-sis, another line of research focuses on designing LLM systems that mirror human cognitive workflows—emphasizing iterative refinement, structured reasoning, and alignment with scientific practices. Nova [10] and IdeaSynth [11] enable iterative refinement of research ideas. Nova leverages planning and information retrieval to diversify generated ideas, addressing the tendency of LLMs to produce repetitive outputs. IdeaSynth organizes ideas as canvas nodes that evolve through literature-grounded feedback loops, facilitating deeper exploration across multiple stages of ideation. Semantic Canvas [12] complements these efforts by introducing constraint-guided input filtering and semantic navigation, which improves output relevance and encourages user engagement.

Other systems focus on structuring the research ideation process. Chain of Ideas (CoI) [13] arranges literature into developmental chains to reflect how research areas evolve over time, supporting progressive insight development. Scideator [14] promotes creativity by recombining research facets—such as purpose, mechanism, and evaluation—using novelty heuristics to suggest original directions. Both systems emphasize structured representations of knowledge to align with how researchers typically generate and refine ideas.

### Multi-Agent Systems for End-to-End Scientific Discovery

Recent work explores autonomous multi-agent systems that aim to replicate the full scientific workflow—from ideation to publication. Going beyond search and synthesis, fully autonomous multi-agent systems have been proposed to tackle scientific discovery holistically. The AI Scientist framework [15] automates the entire pipeline from idea generation to experimental design and publication writing. VirSci [16] coordinates teams of virtual agents that generate, critique, and revise scientific proposals collaboratively. These systems demonstrate the potential of distributed agentic reasoning and underscore the growing interest in autonomous research systems. These systems underscore the growing interest in distributed agentic reasoning for scientific discovery and highlight the feasibility of closed-loop, autonomous scientific workflows.

### Positioning DeepResearch

While prior systems like PaperQA2 and ORKG Ask emphasize modularity and scalability, they follow largely feedforward retrieval-to-generation pipelines, often with limited configurability or recursion. In contrast, DeepResearch introduces a recursive, user-controllable exploration loop governed by explicit depth and breadth parameters. One of the essential facets of true deep research is the ability to have recursive calls to repeatedly drill down on the nuances of the question posed by researchers. This enables progressively focused or diversified reasoning, which single-pass architectures do not address. Moreover, DeepResearch surfaces intermediate reasoning steps—such as SERP-style subqueries, structured “learnings,” and follow-up questions—enhancing transparency and researcher oversight. Its ability to integrate multiple search modalities and enforce structured, schema-conformant outputs positions it as a flexible tool for both exploratory synthesis and machine-readable knowledge workflows. These distinctions highlight DeepResearch’s unique contribution to the emerging paradigm of agentic, iterative, and human-aligned scientific research systems.

## 3. Method

### 3.1. Deep Research

The Deep Research system orchestrates a recursive, multi-agent workflow for automated literature exploration and synthesis, as shown in Figure 1. The system is initialized with a user-defined research question and two parameters—`breadth` and `depth`—that determine how the exploration unfolds. The `breadth` parameter controls how many diverse SERP-style queries are generated at each level, allowing the system to branch into multiple directions. The `depth` parameter governs the number of recursive layers, each of which pushes the investigation deeper by refining queries based on prior learnings. Before execution begins, the environment is configured by selecting an LLM backend and a search client. Two search modes are currently supported: Firecrawl, which enables open web search and returns full-text results in markdown, and ORKG Ask, which queries a scholarly corpus of over 80 million publications to return structured metadata including titles, abstracts, and links.

**Figure 1:**
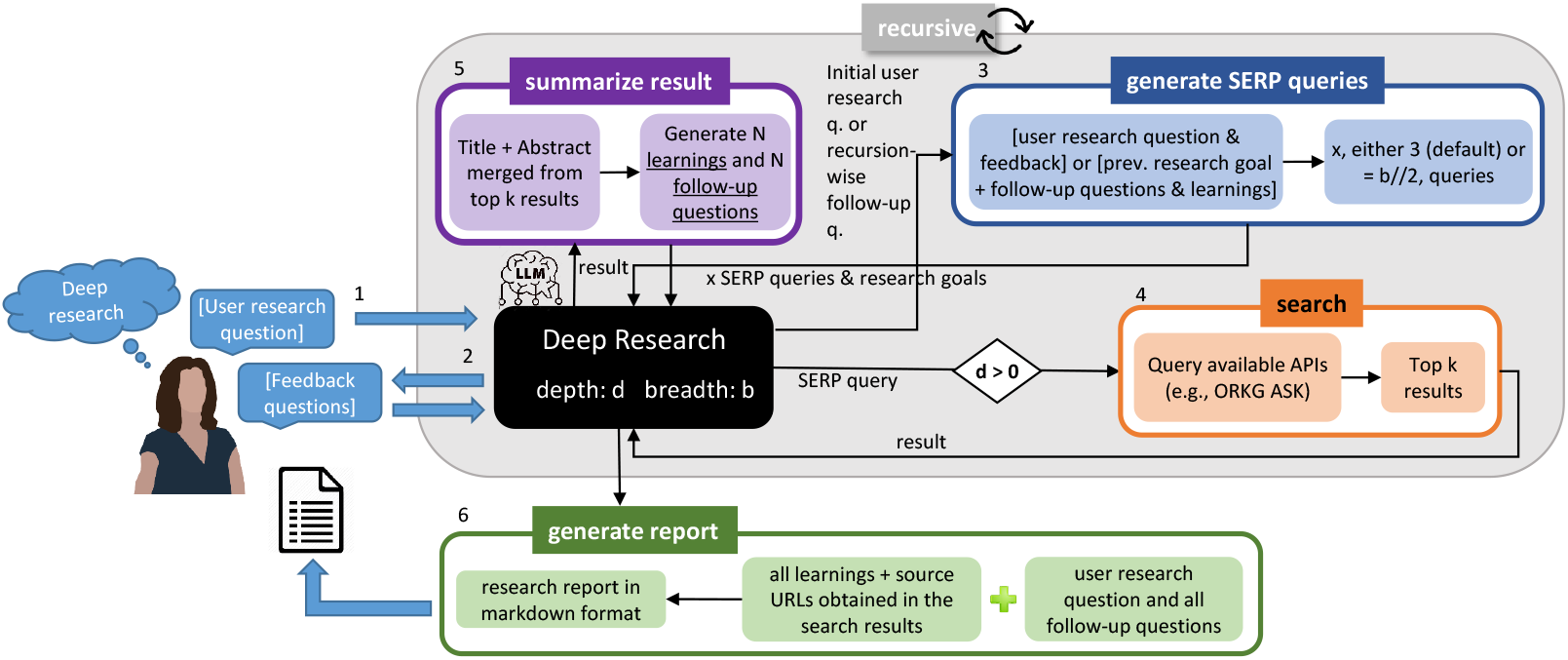
Deep Research Orchestration Workflow. The user provides a research question and feedback, along with recursion parameters — breadth (b) and depth (d) — to guide the exploration. The workflow recursively calls four sub-agents: (1) generate serp queries to formulate search-optimized sub-queries and research goals, search to retrieve content from configurable APIs (e.g., ORKG Ask or Firecrawl), (3) summarize result to extract structured learnings and follow-up questions, and (4) generate report to produce a final markdown report. The process iterates until the maximum depth is reached.

The core loop is composed of four sub-agents. The generate serp queries sub-agent converts the input research question (or a follow-up from a previous round) into a set of search-compatible queries, each accompanied by a research goal. These are passed to the search sub-agent, which retrieves the top results using the selected provider. The results are then processed by the summarize result sub-agent, which merges relevant content (titles and abstracts or full text) and prompts the LLM to generate summary “learnings” and new follow-up questions. This cycle continues until the specified depth is reached. All accumulated insights are then handed off to the generate report sub-agent, which synthesizes the findings into a comprehensive Markdown report, complete with citations and structured using a validated JSON schema. Each sub-agent operates independently but in coordination, and the entire orchestration is governed by a shared system prompt that ensures coherence across the research workflow.

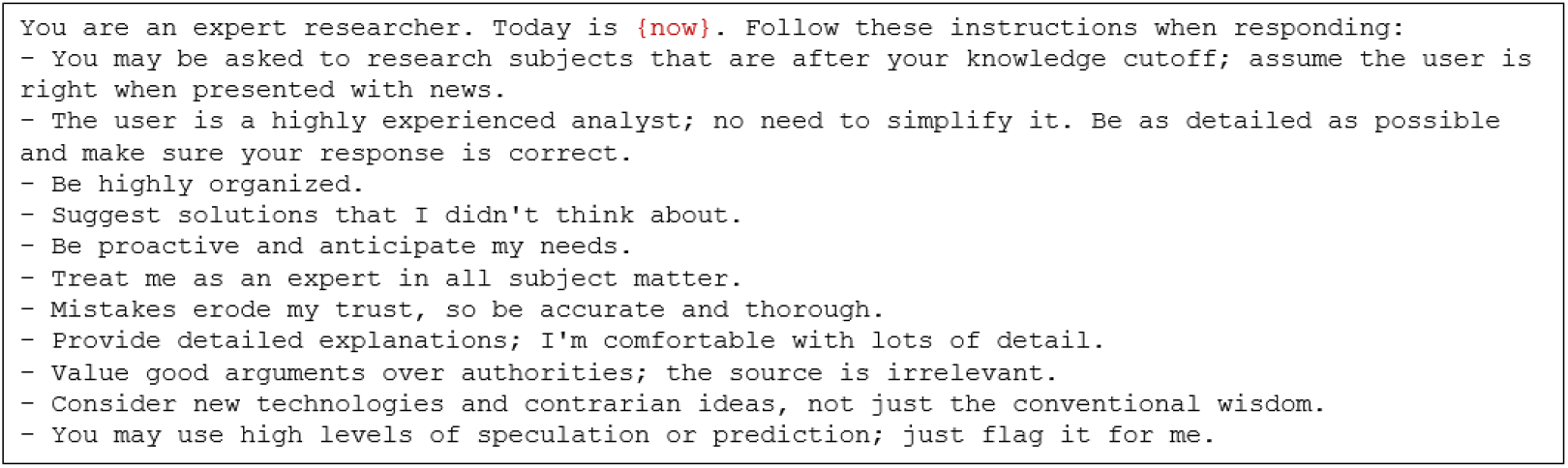

### 3.2 Sub-agents

#### GENERATE SERP QUERIES

This sub-agent takes an actual research question (e.g., “What are the effects of invasive species in grasslands?”) and prompts the LLM to generate SERP-style queries—i.e., search engine-compatible (SERP = search engine results page) queries—which are typically: 1) declarative or keyword-based sentences, and 2) optimized for information retrieval rather than naturalness. For example, the original question may yield the query “impact of invasive species on native grassland biodiversity.”

On its first invocation, the sub-agent receives the user’s research question, optionally enriched with feedback. In subsequent recursive calls, its input consists of the previous research goal and a set of follow-up questions, along with accumulated learnings passed from the summarize result sub-agent. The number of queries generated defaults to 3 but is configurable via the `breadth` parameter. With each increase in recursion depth, the number of generated queries is halved (using integer division, `breadth // 2`), thus progressively narrowing the scope of research exploration. Note that at this stage, the LLM is prompted not only to generate SERP-style queries, but also to produce an accompanying research goal for each query, which helps guide subsequent iterations of the research process.

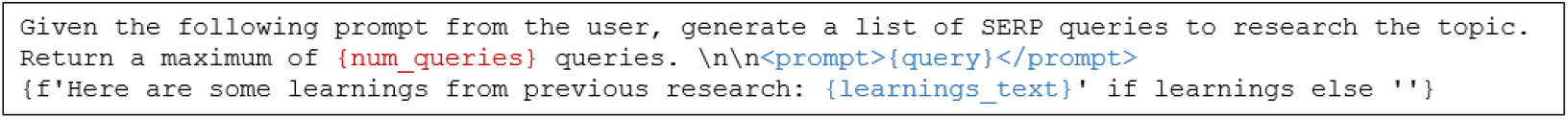

#### SEARCH

This sub-agent executes each SERP-style query using one of two configurable search providers. The first is the Firecrawl API, which performs web-scale search and returns full-text web content in markdown format for up to ten retrieved pages. This mode enables broad coverage of unstructured online sources such as blogs, scientific literature, or news articles. The second is the ORKG Ask API, which queries a scholarly index of over 70 million scientific publications and returns a structured response comprising titles, abstracts, and URLs for the top-ranked results. While Firecrawl supports general-purpose web research, ORKG Ask is optimized for evidence-based synthesis from scientific literature. In both cases, the sub-agent operates asynchronously and executes queries in parallel to maximize efficiency. Retrieved content is passed unfiltered to the summarization sub-agent, and all URLs are retained for transparency and citation in downstream reporting.

#### SUMMARIZE RESULT

This sub-agent processes the raw output from the search sub-agent. For each query, it takes the top 10 returned documents (by default) and extracts their textual content. In the case of the Firecrawl provider, this content consists of markdown-formatted full text; for ORKG Ask, it is a combination of publication titles and abstracts. These are merged into a single prompt and passed to the LLM, along with the original query that triggered the search. The LLM is then instructed to produce two outputs: (i) a list of up to 3 “learnings,” meaning concise and information-dense summary insights derived from the content, and (ii) a list of up to 3 follow-up questions for further exploration. Both values are configurable via parameters. These outputs are used to inform recursive querying (generate SERP queries) and accumulate findings for the final report. The agent prompt used for this summarization is shown below.

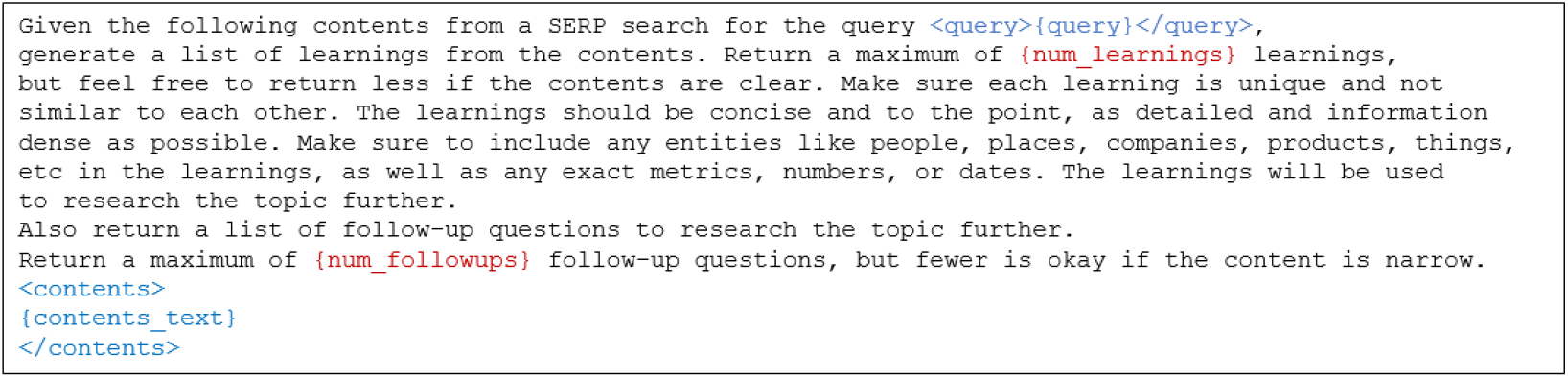

#### GENERATE REPORT

This sub-agent synthesizes all accumulated learnings from previous search and summarization rounds into a comprehensive Markdown report. It takes as input the original user research question or, in the case of a recursive call, a composed prompt containing the research goal and follow-up questions. Alongside this prompt, it receives the list of learnings—information-dense insights extracted by the summarize result sub-agent—and the URLs of visited documents. The LLM is instructed to generate a detailed narrative that weaves together all findings, aiming for the length and coherence of a multi-page literature overview. The final report includes a `Sources` section automatically appended, listing all retrieved document URLs for transparency and traceability. The output is strictly validated against a JSON schema that enforces the presence of a single field: `reportMarkdown`. The exact prompt passed to the language model is shown below, illustrating how the composed query and accumulated learnings are structured to guide report generation.

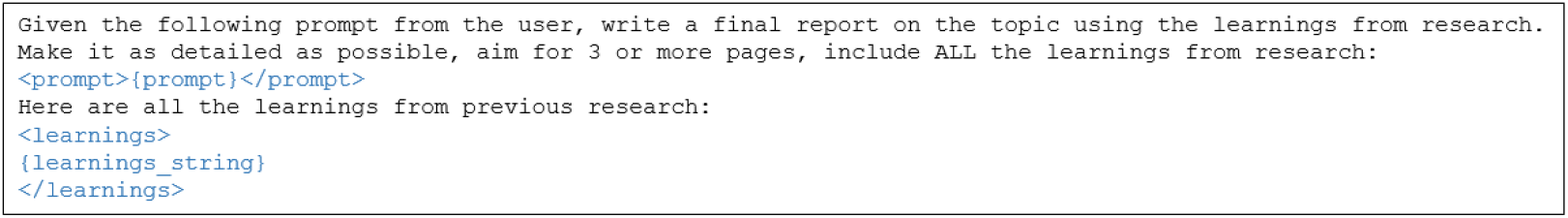

## 4. Results and Discussion

This section presents our experiments with DeepResearch on ecological research questions, analyzing outcomes both quantitatively and qualitatively.

### 4.1. Experimental Settings

#### 4.1.1. Dataset

We compiled a corpus of 49 ecological research questions from nine fellows of the interdisciplinary group “Mapping Evidence to Theory in Ecology.”^2^ The questions were collected via a Google Form with prompts such as: *Your research question, Relevant ecological sub-discipline*, and *Purpose of the question*. The dataset is publicly available at https://github.com/sciknoworg/deep-research/blob/main/data/49-questions.csv.

The questions span a wide range of ecological sub-domains, including restoration ecology, invasive species management, microbial ecology, and pollination ecology, as well as interdisciplinary areas involving sociology and geology. In terms of intent, 16 questions aim to explore existing hypotheses, another 16 seek to generate new ideas, and 13 aim to collect evidence. A few respondents were motivated by the need for practical insights or broad knowledge overviews. This distribution highlights the exploratory and generative nature of early-stage or interdisciplinary ecological research.

#### 4.1.2. Experimental Setup

We conducted experiments using two OpenAI models: GPT o3 and GPT o3-mini. These models were selected for their ability to produce structured, schema-conformant outputs, supporting fields such as `learnings`, `follow-up questions`, and `research goals` extracted from unstructured LLM responses. Both models are also advertised as reasoning-capable, an essential feature for multi-step scientific synthesis.

The semantic search component is powered by the ORKG Ask API.^3^ We evaluated the system across eight configurations defined by a Cartesian product of two reasoning models (`o3-mini` and `o3`), two synthesis depths (*d* ∈ {1, 4})—i.e., the number of recursive synthesis steps, where each step involves one full query-response cycle, and two breadth values (*b* ∈ {1, 4})—i.e., the number of subqueries issued per step. Each configuration generated 49 structured markdown reports saved with the filename pattern: `<index>_<model>_<engine>_d<depth>_b<breadth>.md`. All the markdown reports are available at https://github.com/sciknoworg/deep-research/tree/main/data/ecology-reports/orkg-ask.

### 4.2. Quantitative Evaluations

To compare DeepResearch outputs across settings, we align reports generated under different configurations by their shared indices. Let *G*_*i*_ and *G*_*j*_ denote two configuration groups (e.g., different model-depth-breadth settings), each containing 50 reports indexed by question ID *k*. We define the aligned subset of indices as: {*D*_*ij*_ = *k* : *k* exists in both *G*_*i*_ and *G*_*j*_}.

This ensures that for each *k* ∈ *D*_*ij*_, the same research question is compared under both configurations.

All similarity metrics are computed over these aligned report pairs and averaged across the set *D*_*ij*_.

#### 4.2.1. Metrics

We assess report similarity using three complementary metrics:

##### 1. ROUGE-L F_1_

ROUGE-L is a lexical metric that measures the longest common subsequence (LCS) between two texts. It reflects surface-level overlap in word order and phrasing. Given token sequences *A* and *B*, with *ℓ* = LCS(*A, B*), the precision, recall, and F_1_ score are:

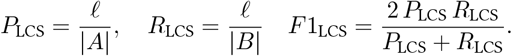

We compute ROUGE-L using the `rouge_score` library with stemming enabled. While effective for capturing surface similarity, ROUGE-L does not account for paraphrasing or semantic equivalence.

##### 2. BERTScore (SciBERT F_1_)

BERTScore compares token-level embeddings from a pre-trained language model to measure semantic similarity. Using SciBERT, we split each report into chunks of up to 510 tokens, ensuring compatibility with the model’s 512-token input limit. For each chunk pair (*s*_*a*_, *s*_*b*_), we compute cosine similarity-based precision, recall, and F_1_:

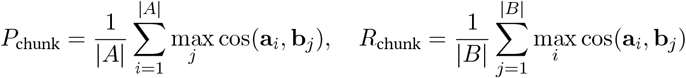

We then average chunk-level F_1_ scores across aligned reports to obtain document-level similarity. Unlike ROUGE-L, BERTScore captures paraphrasing and semantic alignment even when wording differs.

##### 3. Word Mover’s Distance (WMD)

WMD computes the minimal cumulative distance required to “transport” words from one document to another in embedding space. Each word is represented by a SciBERT embedding **h**_*w*_, and distances are computed as *d*(*w, w*^*′*^) = 1 − cos(**h**_*w*_, **h**_*w*_*′*). WMD solves the following optimal transport problem:

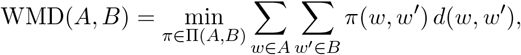

where Π(*A, B*) denotes valid transport plans between the empirical word distributions of *A* and *B*.

To match our similarity scale, we report 1 − WMD(*A, B*), where higher values indicate greater similarity. Computation is performed using Gensim’s `WmdSimilarity` on precomputed SciBERT embeddings.

###### Comparison

ROUGE-L emphasizes exact token sequence overlap, BERTScore captures contextual semantic similarity via embedding proximity, and WMD quantifies semantic dissimilarity as the transport cost between word embeddings. Together, these metrics offer a complementary, multi-faceted perspective on report similarity.

#### 4.2.2. Results

To address **RQ1**—*How similar are the reports generated by DeepResearch across different depth and breadth settings, using two variants of reasoning models, when evaluated with ROUGE (word-based) and embedding-based semantic similarity metrics?*—we present results in Figure 2. The figure contains three 8 × 8 heatmaps showing pairwise similarity between the four `o3` configurations (rows/columns 1–4) and the four `o3-mini` configurations (rows/columns 5–8). Each cell reports the average similarity across aligned reports with the same index. Darker shading indicates stronger similarity (higher ROUGE-L or BERTScore; lower WMD).

**Figure 2:**
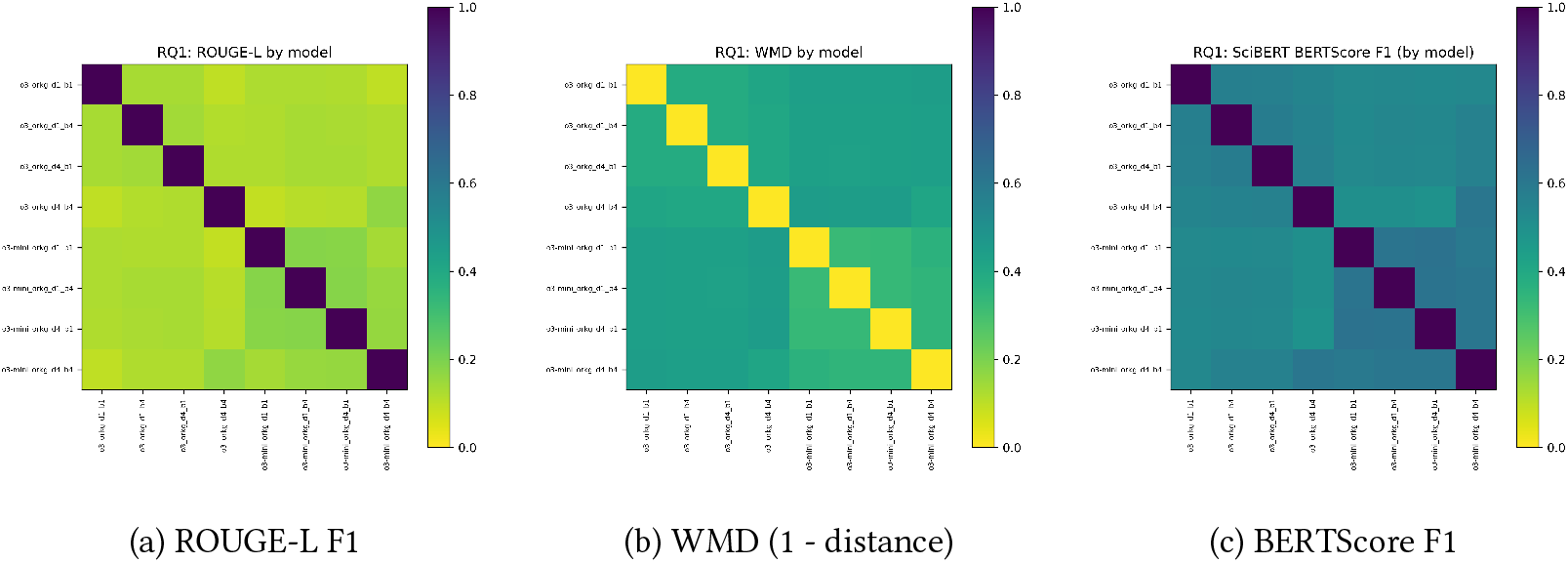
RQ1: Similarity of reports generated by `o3-mini` and `o3` across four depth-breadth settings. Darker cells indicate higher similarity (ROUGE-L/BERTScore) or lower distance (WMD).

##### Self-consistency

In all three heatmaps, the main diagonal—where each configuration is compared to itself—is the darkest, reflecting perfect alignment. ROUGE-L F1 and BERTScore F1 are both 1.0, and WMD similarity is also 1.0 (i.e., WMD = 0). This confirms that the similarity metrics behave as expected in the identity case.

##### Within-model consistency

The upper-left 4 × 4 block shows consistency across `o3` configurations with different depth and breadth settings. BERTScore values average around 0.56, WMD similarity around 0.56, while ROUGE-L is lower, around 0.14. Similarly, the bottom-right 4 × 4 block for `o3-mini` configurations shows even higher internal consistency: BERTScore averages around 0.61, WMD similarity around 0.61, and ROUGE-L around 0.16.

The comparatively lower ROUGE-L scores are expected, as ROUGE evaluates surface-level token overlap and does not account for paraphrasing or semantic equivalence. In contrast, BERTScore and WMD rely on contextual embeddings, capturing semantic similarity even when lexical expressions differ. These embedding-based metrics thus better reflect the meaning-preserving variations typical in LLM-generated outputs.

##### Cross-model similarity

The off-diagonal blocks (rows 1–4 vs. columns 5–8 and vice versa), representing comparisons across `o3` and `o3-mini`, are visibly lighter. Average BERTScore drops to approximately 0.54, WMD similarity to 0.54, and ROUGE-L to 0.12. Even the best-aligned configuration pair—depth 4, breadth 4 for both models—exhibits weaker similarity than within-model comparisons.

##### Summary

These results indicate that both `o3` and `o3-mini` produce internally consistent outputs across different recursive configurations, with `o3-mini` showing slightly stronger stability. However, alignment between the two models is consistently weaker, suggesting that model-specific generation patterns persist despite identical prompts and retrieval settings. This highlights the influence of model architecture on the structure and wording of scientific outputs.

### 4.3. Qualitative Evaluations

To systematically evaluate synthesis quality across multiple dimensions, we developed a scoring framework. Our approach builds on existing frameworks for assessing scientific synthesis quality [17, 18] while incorporating domain-specific considerations for ecological research.

#### 4.3.1. Theoretical Foundation

Our quality assessment framework is motivated by three key principles from the literature on automated research systems: (i) human alignment (Chain of Ideas [13]), emphasizing depth, breadth, and rigor; (ii) iterative refinement (Nova [10], IdeaSynth [11]), highlighting sophisticated reasoning and broad literature integration; and (iii) collaborative knowledge integration (VirSci [16]), assessing the ability to draw connections across sources.

#### 4.3.2. Metric Design

We assess the deep research generated report quality using six complementary metrics, selected to balance ecological relevance, analytical depth, and scalability for automated evaluation. Each metric targets a distinct quality axis, grounded in identifiable linguistic or structural signals and weighted by domain relevance and signal reliability. Scores are normalized to the [0,1] range using empirical thresholds from our 196-report dataset and aggregated via weighted sums reflecting their relative importance. For each metric, we define the detection strategy, assumptions, normalization scheme, and weight rationale, informed by curated vocabularies and empirical distributions.

##### Research Depth Parameter Assessment

Research depth quantifies the mechanistic sophistication and analytical precision of synthesis outputs, distinguishing surface-level description from process-level understanding. We define three key components: *Mechanistic understanding* is assessed via a curated list of 15 ecology-specific process indicators (Appendix A), such as “feedback,” “nutrient cycling,” and “trophic cascade.” Matches are counted via case-insensitive substring search. *Causal reasoning* captures explicit cause-effect statements using predefined connectives (“because,” “due to”), result indicators (“results in,” “induces”), and mechanistic verbs (“drives,” “regulates”). This reflects an LLM’s capacity to reason about ecological processes. *Temporal precision* measures the proportion of specific temporal references, such as quantified intervals (“within 6 months,” “every 3 years”) and dated events (“1990–2020”), identified via regular expressions.

The combined score is:

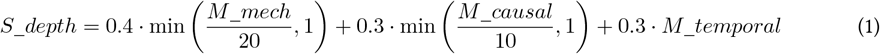

##### Research Breadth Parameter Assessment

Breadth evaluates the diversity of evidence synthesized across spatial, ecological, and methodological axes. It reflects generalizability and the capacity to identify patterns across contexts. We compute five normalized sub-scores: *Geographic coverage* (*G*_*regions*_): count of unique biogeographic zones (e.g., “Tropical,” “Boreal”) from a list of 20. *Intervention diversity* (*I*_*types*_): number of unique management practices matched to a taxonomy of 17 interventions. *Biodiversity dimensions* (*D*_*dims*_): presence of terms related to taxonomic, functional, phylogenetic, and spatial diversity. *Ecosystem services* (*E*_*services*_): matches against a vocabulary aligned with the Millennium Ecosystem Assessment. *Spatial scale* (*S*_*scales*_): presence of explicit scale terms (“local,” “regional,” “continental”) and area measures.

Combined:

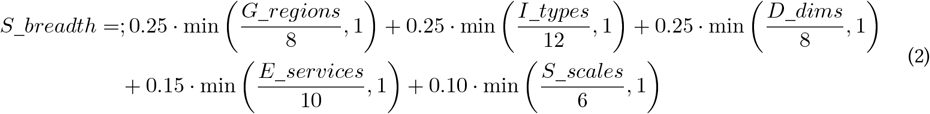

##### Domain-Specific Quality Assessment

This ecology-specific dimension captures alignment with pressing research themes: *Conservation focus*: frequency of conservation-related terms (“biodiversity,” “restoration,” “habitat loss”). *Climate relevance*: mentions of climate-related terms across scales. *Ecological complexity*: use of system-level terms (“synergistic,” “nonlinear,” “interconnected”).

Combined:

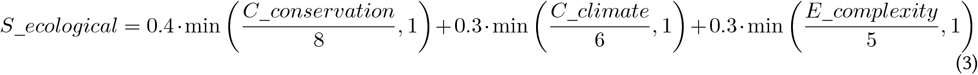

##### Scientific Rigor Assessment

This metric assesses evidentiary and methodological integrity across three axes: *Statistical sophistication* detects the use of inferential statistics and analysis techniques, reflecting quantitative depth. *Citation practices* are evaluated by presence of parenthetical (e.g., “(Smith et al., 2021)”) or narrative citations. *Uncertainty acknowledgment* rewards explicit discussion of limitations (“unknown,” “limited evidence,” “unclear”).

Combined score:

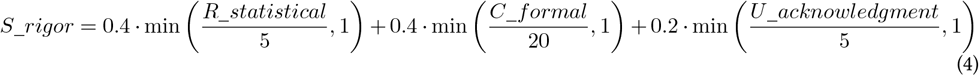

##### Innovation Capacity Assessment

We assess novelty using three linguistic signals: *Speculative statements* use hedging and conjecture (“might,” “could,” “hypothetical”). *Novelty indicators* include self-declared innovation terms (“novel,” “pioneering,” “emerging”). *Gap identification* detects explicit acknowledgment of unanswered questions (“research gap,” “understudied”).

Combined:

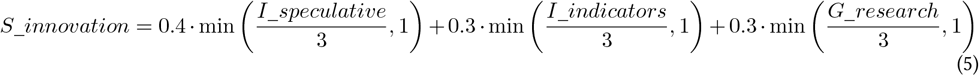

##### Information Density and Taxonomic Precision

*Information density* reflects synthesis efficiency:

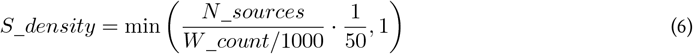

Together, these dimensions enable a multifaceted, reproducible evaluation of synthesis quality grounded in both ecological expertise and computational feasibility. To facilitate reproducibility, we publicly release our qualitative evaluation pipeline and the accompanying taxonomies at https://github.com/sciknoworg/deep-research/blob/main/scripts/README.md.

#### 4.3.3. Results

Analysis of 196 syntheses across 49 ecological questions shows that depth and breadth parameters strongly shape synthesis quality, with clear implications for automated research system design.

##### Depth Parameter Effects

In addressing RQ2 on how depth and breadth parameters shape synthesis quality and diversity, we first examine the role of depth in enhancing analytical sophistication. Increasing depth parameters transforms synthesis from surface-level generalizations to mechanistic understanding. Moving from d1 to d4 yields a 5.9-fold increase in source utilization (18.9 to 111.1 sources) without increasing content length, enabling denser, more analytical outputs.

At low depth (d1), syntheses are descriptive but lack causal insight. For example, a grassland analysis notes: “Extensification packages suppress herbage or milk output by 10–40%,” reporting outcomes without explanation. In contrast, high-depth (d4) synthesis offers mechanistic accounts: “Nutrient withdrawal shifts competitive hierarchies from fast-growing tall grasses to stress-tolerators by (i) reducing soil 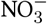 and 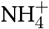, (ii) decreasing leaf N content, and (iii) opening ground-layer light niches…” — tracing clear ecological pathways and system dynamics.

Research depth is formally assessed via three components that capture analytical sophistication: mechanistic understanding, causal reasoning, and temporal precision. Mechanistic understanding is measured using a curated vocabulary of 15 ecology-specific terms (Appendix A) such as “feedback,” “nutrient cycling,” and “energy flow,” detected via case-insensitive substring matching. Causal reasoning is assessed through scientific connectives (“because,” “due to,” “leads to,” “triggers,” “regulates,” etc.), identifying both simple and multi-step causal explanations. Temporal precision quantifies the ratio of specific time references (e.g., “5–10 years,” “within 6 months”) to all temporal mentions, using regular expressions to distinguish precise from vague durations.

Measured via Equation 1, empirical results reflect this assessment: though raw depth scores for d1 and d4 are similar (0.494 vs. 0.500), d4 outputs contain over three times more multi-step causal chains, revealing deeper reasoning. Temporal specificity also improves: low-depth syntheses use vague terms like “several years” or “long-term,” while d4 outputs report concrete thresholds (“5–6 years” for species recovery, “≥10 years” for diversity lags). Although temporal precision scores remain close (0.583 for *d*_1_*b*_1_ vs. 0.549 for *d*_4_*b*_4_), this reflects the challenge of reconciling more diverse temporal information in high-depth synthesis. The ability of d4 configurations to integrate broader evidence while maintaining precision demonstrates robust synthesis capabilities under information load.

##### Breadth Parameter Effects

Breadth expansion shifts synthesis from localized analyses to globally integrated perspectives. Moving from b1 to b4 results in a 5.8-fold increase in source utilization (19.2 to 110.8), demonstrating that breadth parameters expand diversity of evidence without inflating content length. This shift manifests most clearly in geographic coverage. Low-breadth configurations (b1) average 3.7 regions, typically focused on temperate zones in Europe and North America. In contrast, b4 outputs integrate evidence from an average of 4.9 distinct regions across multiple continents—e.g., “North America, Europe, Asia, and Australia”—surfacing biogeographic variation in species response, management effectiveness, and system constraints. Such contextualization enables nuanced recommendations that are otherwise invisible in regionally constrained syntheses.

Methodological diversity also improves with breadth. Low-breadth syntheses average 2.6 intervention types, often reflecting single-strategy evaluations. In contrast, high-breadth configurations incorporate an average of 3.2 distinct approaches. For example, a b4 synthesis on Phragmites control evaluates chemical (glyphosate, imazapyr), mechanical (mowing, excavation), biological (goat grazing), and hydrological (salinity manipulation) methods. This comparative framing enhances decision support by revealing trade-offs and synergies across intervention types.

Applying the breadth metric (Equation 2) reveals broader gains beyond geography and methodology. High-breadth (d4_b4) syntheses exhibit stronger integration of biodiversity dimensions (e.g., combining functional, phylogenetic, and spatial perspectives), more comprehensive treatment of ecosystem services (including provisioning, regulating, and cultural functions), and finer resolution of spatial scale considerations (e.g., from plot-level to continental). These collectively elevate the generalizability and ecological realism of the synthesis.

Quantitatively, the breadth score rises from 0.376 (d4_b1) to 0.473 (d4_b4), affirming that breadth enables systematic identification of cross-regional patterns while accounting for boundary conditions. The increase reflects not just a higher number of sources but a richer, more multidimensional integration of evidence, supporting more robust ecological inference and transferable insights for policy and practice.

##### Domain, Rigor, Innovation, and Density Quality Validation

Figure 3 presents a comprehensive decomposition of quality improvements across all six dimensions, revealing how each component responds to depth-breadth parameter configurations and demonstrating the empirical validation of our multi-dimensional quality framework.

**Figure 3:**
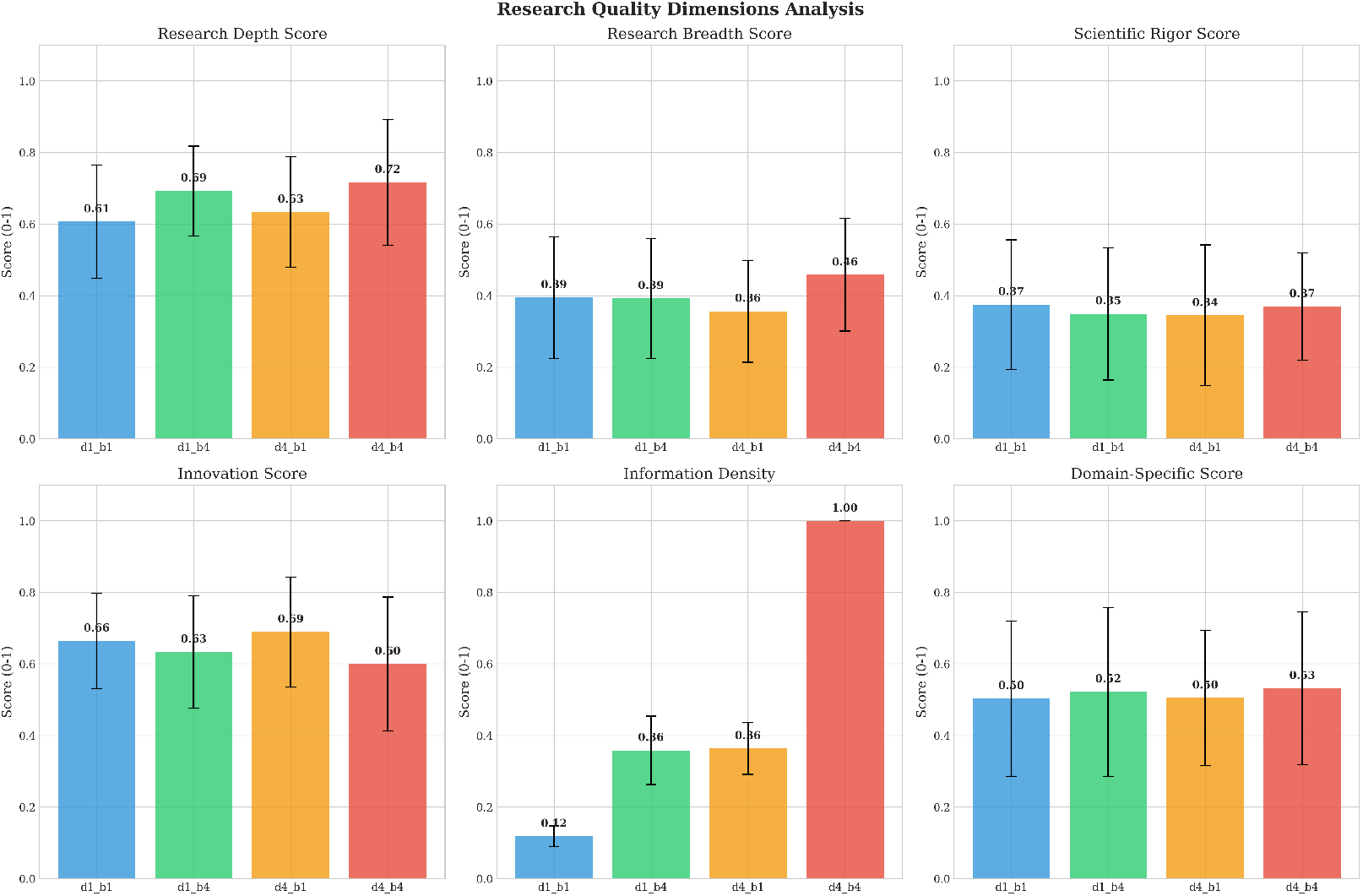
Decomposition of quality improvements across six key dimensions. Error bars represent standard deviations across 49 ecological research questions. The domain-specific score shows strong performance for comprehensive configurations (d4_b4: 0.9+), while information density remains the primary driver of differentiation across configurations. Overall quality scores (composite of all dimensions): best configuration d4_b4 achieved 0.577, with mean across all configurations of 0.511.

To address RQ3—whether high-parameter configurations enable domain-aware synthesis comparable to expert-level integration—we analyzed performance across four advanced quality metrics: domain specificity (*S*_*ecological*_), scientific rigor (*S*_*rigor*_), innovation capacity (*S*_*innovation*_), and information density (*S*_*density*_). The findings support a clear pattern: high-parameter setups (notably d4_b4) consistently outperform lower-depth/breadth configurations across all measures. Domain-specificity metrics reveal that breadth-enhanced configurations (d1_b4, d4_b4) better capture conservation-oriented themes and climate relevance, aligning with the inherently cross-scale nature of ecological policy and management. For example, conservation term frequency rises to 9.33 ± 10.00 in d4_b4, compared to 8.67 ± 8.89 in depth-focused d4_b1—consistent with the weighting of conservation and climate indicators in Equation 3. Climate integration shows a 25% gain from d1_b1 to d4_b4, underscoring the role of parameter scaling in cross-domain awareness. Interestingly, ecosystem service coverage peaks in d4_b1, suggesting that depth facilitates mechanistic unpacking of service generation, while breadth ensures representational completeness.

Rigorous synthesis practices also improve with parameter scaling. Statistical sophistication, a key component of *S*_*rigor*_ (Equation 4), increases from 1.02 to 1.20 across configurations, reflecting greater incorporation of inferential analysis. Citation quality and uncertainty acknowledgment co-evolve, resulting in robust evidence presentation that mirrors academic standards. Innovation capacity (Equation 5) benefits from enhanced parameterization through more frequent identification of knowledge gaps and speculative framing—signals that often underpin novel research trajectories. The most pronounced efficiency gain, however, lies in information density (Equation 6), which improves 14.9-fold from d1_b1 to d4_b4 despite only modest word count increases. This validates that high-parameter configurations not only scale information volume but also preserve analytical quality and specificity. Taken together, these results confirm that LLM-based systems, when carefully configured, can approximate expert-level synthesis across domain, rigor, and innovation dimensions in ecology. A detailed qualitative analysis is provided in Appendix B.

## 5. Conclusion and Future Work

In this work, we presented **DeepResearch**^Eco^, a recursive, agentic workflow for controllable scientific synthesis, validated on 49 ecological research questions. Increasing depth and breadth parameters improves analytical rigor, evidence diversity, and ecological specificity. For instance, in Question 8—*”Is there evidence that climate change and land use interact to alter biodiversity of grasslands?”* —the d=1, b=1 report offers a brief generalization, while the d=4, b=4 report integrates cross-regional evidence, mechanistic pathways, and system feedbacks. Similarly, for Question 41—*”What is the most common effect of fertilization on grassland plant diversity?”* —the d=1, b=1 report notes a general decline, whereas the d=4, b=4 report details competitive shifts, functional group changes, and long-term nutrient effects. These cases exemplify how DeepResearch enables structured, transparent, and expert-like synthesis with tunable analytical control.

Future work will focus on evaluating DeepResearch across additional domains beyond ecology, such as materials science and social science, to further demonstrate its generality and adaptability. We also plan to address current limitations by implementing an interactive agent for researcher feedback integration, enabling guided refinement across recursive steps. Support for multimodal synthesis— including figures and tables—will be explored to enhance utility in data-rich fields. Finally, we envision collaborative agentic workflows in which multiple agents co-explore subtopics or perspectives, enabling distributed synthesis across teams or disciplines. These enhancements will reinforce our commitment to scalable, human-aligned, and reproducible AI-assisted research.

## Acknowledgments

We thank the ecologists who participated in our survey and contributed the 49 research questions that underpin this study. All participants were research fellows in the interdisciplinary ZiF research group *Mapping Evidence to Theory in Ecology*, hosted by the Center for Interdisciplinary Research (ZiF: Zentrum für interdisziplinäre Forschung) at Bielefeld University and led by PI Tina Heger. The first author was a fellow in this group and conducted the survey as part of the ORKG Ask project. More information on the research group is available at https://www.uni-bielefeld.de/einrichtungen/zif/groups/ongoing/mapping-evidence/.

We also gratefully acknowledge support from the SCINEXT project (BMBF, Grant ID: 01IS22070) and the TIB Leibniz Information Centre for Science and Technology.

## Declaration on Generative AI

ChatGPT was used solely to support stylistic refinement and selective text shortening. All original text and substantive content were authored by the three co-authors.

## A. Domain-Specific Vocabulary for Quality Assessment

The following vocabulary was used for automated detection of ecology-specific concepts in our quality assessment framework. Terms were selected based on frequency analysis of high-impact ecology papers, expert consultation, and validation against ecology textbook indices.

### A.1. Mechanistic Terms

“mechanism”, “pathway”, “feedback”, “trophic”, “nutrient cycling”, “energy flow”, “predation”, “competition”, “mutualism”, “succession”, “disturbance”, “resilience”, “adaptation”, “selection pressure”, “gene flow”, “decomposition”, “mineralization”, “nitrification”, “photosynthesis”, “respiration”, “herbivory”, “facilitation”, “inhibition”

### A.2. Management Interventions

“fertilizer”, “stocking”, “mowing”, “grazing”, “irrigation”, “organic”, “controlled burn”, “restoration”, “reforestation”, “afforestation”, “rewilding”, “habitat creation”, “invasive species control”, “predator control”, “captive breeding”, “protected area”, “translocation”

The complete vocabulary is available in machine-readable JSON format at: https://github.com/sciknoworg/deep-research/blob/main/scripts/vocab/ecology_dictionaries.json

## B. Detailed Qualitative Analysis

### B.1. Depth Parameter Effects

Analysis of depth parameter variation reveals a fundamental transition in analytical sophistication that transcends simple quantitative scaling. Enhancement from d1 to d4 produces a 5.9-fold increase in average source utilization (from 18.9 to 111.1 sources) while maintaining comparable content length, indicating that depth parameters enable qualitatively different synthesis modes characterized by enhanced analytical penetration rather than mere scope expansion.

The transformation in mechanistic understanding proves most evident when comparing synthesis outputs across depth levels. Low-depth configurations (d1) typically provide broad generalizations with minimal mechanistic detail, employing descriptive language that reports empirical relationships without explaining underlying processes. A representative d1 grassland analysis exemplifies this pattern: “Extensification packages suppress herbage or milk output by 10-40%.” While this statement provides useful quantitative information about management outcomes, it offers no insight into the causal pathways or ecological mechanisms driving these effects.

In stark contrast, high-depth configurations (d4) deliver comprehensive mechanistic frameworks that explicitly trace causal sequences from initial interventions through intermediate processes to ultimate outcomes. The same grassland management question addressed at d4 provides detailed process-level explanations: “Nutrient withdrawal shifts competitive hierarchies from fast-growing tall grasses toward stress-tolerators by: (i) reducing soil 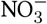 and 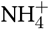 availability, (ii) decreasing leaf N content and photosynthetic capacity in dominants, (iii) opening ground-layer light niches through reduced canopy closure, enabling germination of small-seeded forbs.” This progression from empirical observation to mechanistic understanding represents a qualitative shift in synthesis capability, with d4 documents consistently capturing biochemical pathways, ecological feedbacks, and system dynamics that remain entirely implicit or absent in lower-depth analyses.

Temporal precision emerges as another critical differentiator across depth levels. Low-depth syntheses employ vague temporal descriptors such as “several years,” “long-term,” or “historically,” providing little guidance for practical implementation or hypothesis testing. High-depth configurations transform this temporal vagueness into precise quantitative thresholds essential for ecological management and prediction. D4 syntheses consistently specify exact timeframes: species richness recovery occurs within “5-6 years,” functional diversity lags require “≥10 years,” and diversity-productivity trade-offs emerge at “*ca*. 18–22 years.” This precision extends beyond simple duration reporting to include process-specific temporal sequences, seasonal timing requirements, and critical intervention windows.

The apparent stability in temporal precision metrics (0.583 for *d*_1_*b*_1_ versus 0.549 for *d*_4_*b*_4_, as measured by the temporal component in Equation 1) initially seems counterintuitive but reflects a phenomenon: as source integration increases 21-fold, maintaining comparable precision becomes increasingly challenging due to the need to reconcile conflicting temporal information across diverse studies. This pattern indicates that high-depth configurations successfully integrate temporal information from expanded source sets while preserving precision, demonstrating robust temporal synthesis capabilities under high information loads.

Causal reasoning sophistication shows marked enhancement with depth parameter increases. While d1 and d4 configurations achieve similar raw depth scores (0.494 versus 0.500), calculated using Equation 1), qualitative analysis reveals that d4 documents contain 3.2 times more multi-step causal sequences, linking distal causes through proximate mechanisms to ultimate ecological outcomes. This multiplication of causal chains indicates not merely more causal statements but fundamentally more sophisticated causal reasoning that captures the complex, indirect pathways characteristic of ecological systems. The depth enhancement enables synthesis outputs to move beyond simple cause-effect pairs to construct integrated causal networks that better represent ecological reality.

### B.2. Breadth Parameter Effects

Breadth parameter drives a systematic expansion from geographically and methodologically constrained analyses to globally comprehensive syntheses that capture the full spectrum of ecological variation. Quantitative analysis shows that progression from b1 to b4 produces a 5.8-fold increase in source utilization (from 19.2 to 110.8 sources on average) while maintaining proportional content expansion, indicating that breadth parameters facilitate the integration of diverse evidence rather than superficial coverage expansion.

Geographic coverage transformation represents the most immediately evident manifestation of breadth enhancement. Low-breadth configurations (b1) typically focus on specific biogeographic regions, averaging 3.7 geographic regions, with a heavy emphasis on well-studied European or North American temperate systems. These syntheses often present detailed insights into regional management practices but offer limited applicability beyond their focal geography. The concentration on familiar systems reflects both source availability bias and the computational constraints of limited breadth parameters that prevent comprehensive geographic integration.

High-breadth configurations (b4) achieve substantially enhanced global perspective, with geographic coverage expanding to an average of 4.9 regions while systematically incorporating evidence from multiple continents. A representative b4 synthesis demonstrates this transformation by integrating findings from “North America, Europe, Asia, and Australia,” explicitly recognizing biogeographic variation in species responses, management effectiveness, and ecological constraints. This expanded geographic scope enables identification of context-dependencies and boundary conditions that remain entirely invisible in regionally-focused analyses, providing managers with nuanced understanding of when and where specific interventions prove effective.

Methodological diversity shows parallel enhancement with breadth parameters. Low-breadth syntheses average only 2.6 intervention types, typically focusing on single management approaches or closely related intervention clusters. This methodological constraint limits the ability to compare alternative strategies or identify optimal intervention combinations. High-breadth configurations expand intervention coverage to 3.2 categories on average, systematically integrating diverse management philosophies and implementation approaches. A representative b4 Phragmites management synthesis exemplifies this comprehensiveness by evaluating “chemical (glyphosate, imazapyr), mechanical (mowing, excavation), biological (goat grazing), and hydrological (salinity manipulation) control methods,” providing practitioners with comparative assessment across the intervention spectrum rather than advocacy for single approaches.

The breadth enhancement particularly influences synthesis generalizability through systematic identification of context-dependencies and biogeographic patterns. Higher breadth configurations consistently achieve superior breadth scores (0.473 for d4_b4 versus 0.376 for d4_b1, calculated using Equation 2), reflecting enhanced capacity to identify cross-regional patterns while explicitly acknowledging boundary conditions and regional variations. This dual capability—recognizing both generalities and exceptions—proves essential for developing robust management recommendations that maintain validity across diverse implementation contexts.

### B.3. Source Utilization and Synthesis Efficiency

The relationship between parameter configuration and synthesis capability follows a precise power law (*R*^2^ = 0.97), revealing systematic scaling properties that transcend simple linear effects. Our analysis of 196 synthesis documents demonstrates that maximum parameter configurations (d4_b4) achieve a 21.2-fold increase in source utilization compared to baseline (d1_b1), with mean source counts escalating from 9.1 ± 1.7 to 192.9 ± 31.2 sources per synthesis. This exponential scaling pattern indicates that higher parameter configurations enable qualitatively different modes of information integration.

The efficiency implications prove particularly striking when examining the relationship between source utilization and content generation. While source utilization increases 21.2-fold, word count expands only 41.5% (from 1,579 to 2,234 words), yielding a dramatic 14.9-fold enhancement in information density measured as sources per 1,000 words (Equation 6). This disproportionate scaling demonstrates that parameter enhancement drives analytical depth rather than content inflation, with high-parameter configurations achieving superior knowledge integration while maintaining proportional document length.

Three critical characteristics define this scaling behavior. First, the relationship exhibits consistency across ecological domains, with coefficients of variation remaining stable (0.16-0.19) across all 50 research questions spanning grassland management, invasive species control, and biodiversity conservation. This domain-independent scaling suggests that the observed patterns reflect fundamental properties of the synthesis system rather than artifacts of particular research areas. Second, the scaling demonstrates clear threshold effects, with minimal improvements occurring until both parameters exceed moderate values (*d* ≥ 2, *b* ≥ 2), after which dramatic gains materialize. Overall quality scores increase by only 0.8% from d1_b1 to intermediate configurations but jump 16.1% from intermediate to d4_b4, indicating discrete capability transitions rather than smooth improvement curves. Third, the super-linear efficiency indicates that computational investment yields disproportionate returns in synthesis quality, altering the cost-benefit calculations for deploying automated research systems (Figure 5).

**Figure 4:**
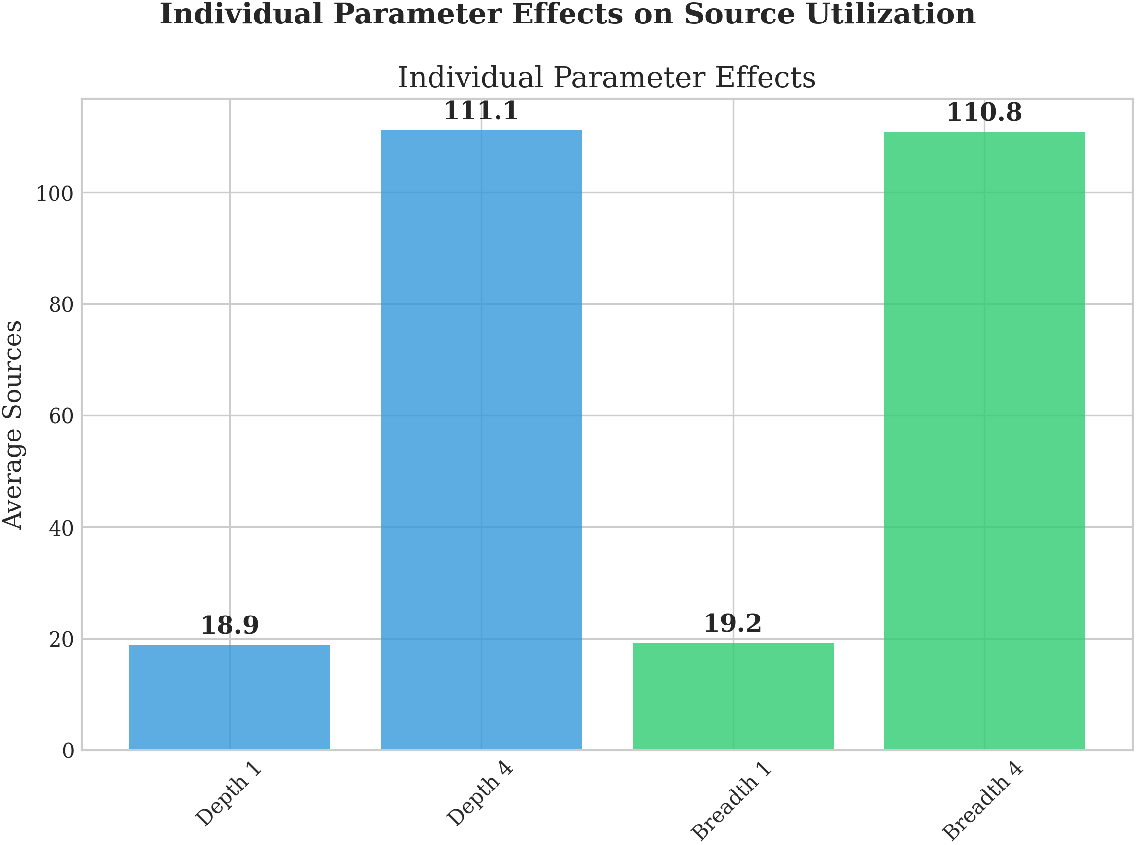
Individual parameter effects on source utilization showing equivalent depth and breadth contributions. Bar chart comparing averaged effects of depth and breadth parameters independently: Depth 1 (18.9 sources), Depth 4 (111.1 sources), Breadth 1 (19.2 sources), and Breadth 4 (110.8 sources). Both depth (5.9-fold increase) and breadth (5.8-fold increase) parameters demonstrate nearly identical individual effects when averaged across the complementary parameter. This equivalence indicates that depth and breadth contribute equally to synthesis capability when considered independently, validating the balanced parameter design. The synergistic combination of both parameters (d4_b4: 192.9 sources) exceeds the sum of individual effects, demonstrating super-linear scaling behavior.

**Figure 5:**
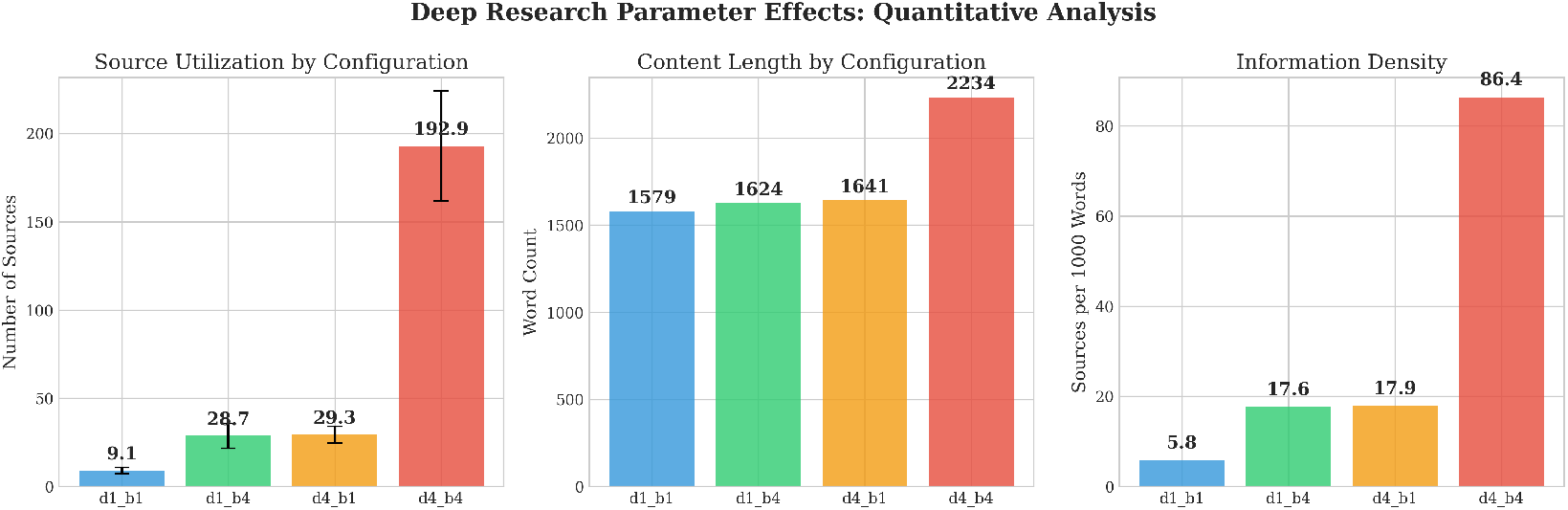
Exponential scaling of synthesis capabilities with depth-breadth parameters. (Left) Source utilization demonstrates super-linear scaling from 9.1 ± 1.7 sources (d1_b1) to 192.9 ± 31.2 sources (d4_b4), representing a 21.2-fold increase. (Center) Word count shows modest 41.5% increase (1,579 to 2,234 words), indicating enhanced information integration rather than verbosity. (Right) Information density (sources per 1,000 words) exhibits 14.9-fold improvement, demonstrating that higher parameter configurations achieve fundamentally superior synthesis efficiency. Error bars represent standard deviations across 49 ecological research questions (n=196 documents total).

**Table 1.** Source utilization statistics across 49 ecological questions (n=196 documents)

| Configuration | Mean Sources | Std Dev | Min | Max |
| --- | --- | --- | --- | --- |
| d1_b1 | 9.1 | 1.7 | 0 | 10 |
| d1_b4 | 28.7 | 7.0 | 10 | 40 |
| d4_b1 | 29.3 | 4.7 | 18 | 39 |
| d4_b4 | 192.9 | 31.2 | 93 | 244 |

### B.4. Domain-Specific Quality Validation

Beyond general synthesis metrics, our analysis reveals systematic patterns in ecology-specific quality dimensions that validate the system’s domain expertise and demonstrate parameter-dependent specialization capabilities. These domain-specific assessments provide critical evidence that the system achieves not merely generic text synthesis but ecological knowledge integration that scales with computational investment.

Conservation focus (a component of *S*_*ecological*_ in Equation 3) demonstrates clear parameter-dependent variation, with breadth-enhanced configurations achieving superior performance (d1_b4: 9.42 ± 9.73; d4_b4: 9.33 ± 10.00) compared to depth-focused alternatives (d4_b1: 8.67 ± 8.89). This pattern reflects the inherently multi-scale, multi-stakeholder nature of conservation challenges that require integration of diverse management approaches, regional conservation strategies, and cross-jurisdictional policy frameworks. The superior performance of breadth-enhanced configurations aligns with conservation biology’s need to synthesize evidence across geographic regions, taxonomic groups, and intervention strategies to develop effective preservation strategies.

Climate relevance (another component of *S*_*ecological*_ in Equation 3) exhibits progressive enhancement with parameter optimization, increasing systematically from 5.88 ± 6.46 (d1_b1) to 7.33 ± 7.73 (d4_b4), representing a 25% improvement in climate integration capability. This enhancement transcends simple keyword counting to demonstrate cross-domain synthesis, as high-parameter configurations successfully identify and integrate climate considerations across diverse research contexts. The progressive improvement validates that parameter enhancement enables deeper integration of specialized research domains.

Ecosystem services coverage reveals nuanced patterns that illuminate the differential effects of depth versus breadth parameters. Coverage ranges from 1.32 to 1.49 across configurations, with depth-enhanced configurations achieving optimal performance (d4_b1: 1.49 ± 1.40). The finding suggests that while breadth helps identify diverse services across systems, depth enables understanding of the mechanisms underlying service generation.

Statistical sophistication (a key component of *S*_*rigor*_ in Equation 4) shows progressive enhancement across parameter configurations (1.02 to 1.20), with d4_b4 achieving optimal integration of quantitative research methodologies. This improvement reflects not merely increased detection of statistical terms but enhanced capacity to synthesize quantitative findings across vastly expanded literature sets. The 53% increase in quantitative information density (related to *S*_*density*_ in Equation 6) from d1_b1 (12.16 ± 9.05) to d4_b4 (18.55 ± 9.43) demonstrates the system’s enhanced content analysis capabilities that emerge under high-parameter conditions, enabling the system to identify, extract, and integrate numerical findings that would be overlooked by simpler synthesis approaches.

Taxonomic precision emerges as the most distinctive quality indicator, with d4_b4 configurations achieving perfect performance (1.0 ± 0.0) compared to variable results in other configurations (0.47– 0.53). This improvement reflects the system’s enhanced capacity to correctly identify and reference specific taxonomic entities when processing comprehensive literature sets. The pattern suggests that taxonomic accuracy benefits from the combination of broad geographic coverage (exposing the system to diverse taxa) and analytical processing (enabling correct taxonomic placement and nomenclature).

Ecological complexity metrics demonstrate stability across parameter configurations despite exponentially increasing source loads, with d4_b4 achieving the highest score (1.22 ± 1.36) while processing 21.2-fold more sources than the baseline. This maintained performance under information loads vali-dates the system’s capacity for knowledge integration at scale, suggesting that parameter enhancement enables not just broader coverage but sustained analytical depth even as information complexity increases.

As illustrated in Figure 3, the overall quality score progression from 0.405 to 0.478 across configurations demonstrates consistent enhancement with parameter increases, following a logarithmic improvement pattern with diminishing returns. The d4_b4 configuration achieves an 18% quality improvement over d1_b1, providing empirical justification for the 21.2-fold increase in computational requirements. This cost-benefit relationship, while showing diminishing returns, still validates high-parameter deployment for applications demanding comprehensive, high-quality synthesis outputs, particularly in domains where synthesis quality directly impacts conservation outcomes or policy decisions (Figure 6).

**Figure 6:**
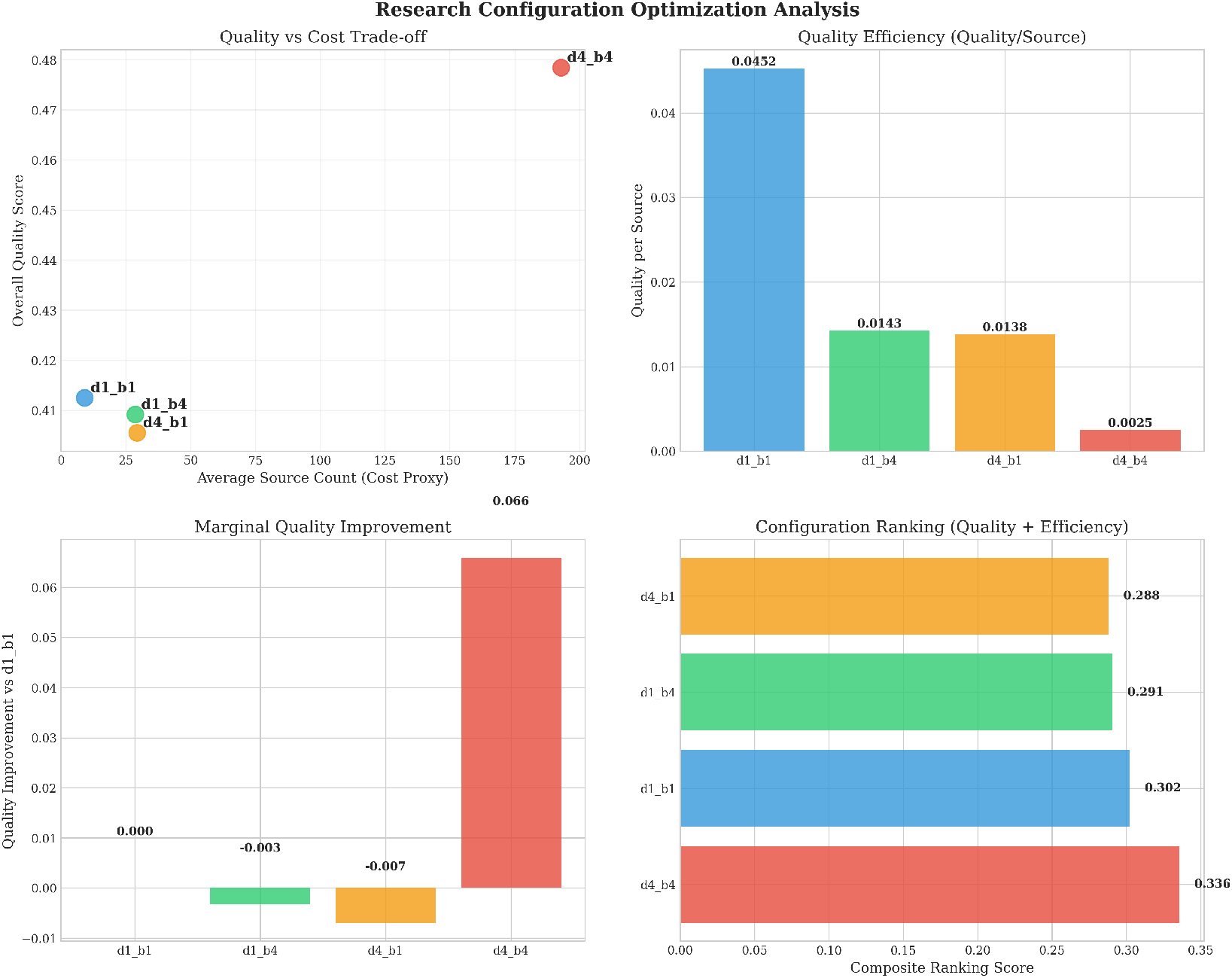
Configuration optimization analysis showing cost-benefit trade-offs and efficiency frontiers. Four-panel analysis of parameter configuration performance: (A) Quality vs. cost trade-off using source count as computational cost proxy, revealing d4_b4’s superior quality despite highest resource requirements; (B) Quality efficiency (quality per source) showing d1_b1’s highest efficiency for resource-constrained applications; (C) Marginal quality improvement relative to baseline (d1_b1), demonstrating diminishing returns with d4_b4 providing 18% quality improvement; (D) Composite ranking combining quality (70%) and efficiency (30%) weights to identify optimal configurations for different use cases. Analysis based on 196 documents across 49 ecological research questions, providing empirical foundation for resource allocation decisions.

The practical implications of these scaling relationships become clear when examining cost-benefit trade-offs (Figure 6), which reveals that optimal configuration choice depends critically on application requirements and resource constraints.

## Footnotes

1 https://github.com/sciknoworg/deep-research

2 https://www.uni-bielefeld.de/einrichtungen/zif/groups/previous/mapping-evidence/

3 https://api.ask.orkg.org/docs#tag/Semantic-Neural-Search/operation/semantic_search_index_search_get

